# Nearly maximal information gain due to time integration in central dogma reactions

**DOI:** 10.1101/2022.01.02.474710

**Authors:** Swarnavo Sarkar, Jayan Rammohan

**Affiliations:** National Institute of Standards and Technology, Gaithersburg, MD 20899, USA

## Abstract

Living cells process information about their environment through the central dogma processes of transcription and translation, which drive the cellular response to stimuli. Here, we study the transfer of information from environmental input to the transcript and protein expression levels. Evaluation of both experimental and analogous simulation data reveals that transcription and translation are not two simple information channels connected in series. Instead, we show that the central dogma reactions often create a time-integrating information channel, where the translation channel receives and integrates multiple outputs from the transcription channel. This information channel model of the central dogma provides new information-theoretic selection criteria for the central dogma rate constants. Using the data for four well-studied species we show that their central dogma rate constants achieve information gain due to time integration while also keeping the loss due to stochasticity in translation relatively low (*<* 0.5 bits).

---

Francis Crick described the central dogma of molecular biology as the unidirectional and sequential flow of information from DNA to RNA to protein through the biochemical reactions of transcription and translation, which is responsible for most cellular response [1, 2]. Although Crick’s definition of “information” differed from the quantitative information-theoretic term [3], it prompts the question: is it possible to rigorously quantify information transfer in cells, from environmental stimuli through transcription and translation? In the past two decades, a combination of experimental and computational progress has demonstrated that information transfer can be quantified in biological systems of varying complexity [4–8]. However, to put Crick’s statement of the central dogma on a quantitative footing, we need to use information theory to examine how does transcription and translation modulate the information about the environment available in cells.

Much of the recent progress in quantifying information transfer in biological systems has been enabled by advances in single-cell measurements, which provides the distribution of cellular responses, in the form of transcript or protein expression levels [9–12]. Previous work has examined information transfer from the environmental input to either the transcript or the protein expression levels, and its dependence on the transcription and translation rate constants [6, 13–15]. However, those studies have not examined information transfer during the elementary and sequential processes of transcription and translation. Instead, they have mostly focused on biological networks [16], cellular decision making [17], or intracellular distribution of information [18]. Though there has been a renewed interest in probing the central dogma in recent years [19, 20], a comprehensive information-theoretic treatment of information transfer from the environment to the cellular response encompassing both transcription and translation is still lacking. Such a treatment of the central dogma will help to explain the central dogma rate constants prevalent in different species and biological pathways, and could inform the design of engineered cellular sensing systems.

Here, we use single-cell measurements and information theory to show that biology achieves nearly maximal information transfer from the environmental input through translation to the protein expression. We find from the analysis of single-cell measurements that the information transfer from the environmental input to the protein expression level is apparently higher than the information transfer from the same environmental input to the transcript expression level. This contradicts an elementary result from information theory that information should be lost through a simple serial connection of information channels [21]. To explain this unexpected gain in information during translation, we develop an information channel model whose information processing properties are functions of the central dogma rate constants. The information channel model highlights two distinct factors that affect the information gain during translation: (1) time integration of the transcript expression, where the amount of signal integration is set by the ratio between the decay rate constants for the transcript and the protein, and (2) the average protein expression level, which is determined by the translation power, *i*.*e*. the ratio between the translation rate constant and the protein decay rate constant. We estimate the translation loss (the amount of information lost during translation) as the difference between the true protein-level information gain and the maximum possible information gain for the same integration time. For a fixed translation power, the information gain from time integration increases with increasing integration time, but the translation loss also increases. By computing the information gain for multiple species, using central dogma rate constants from the literature, we show that the naturally occurring central dogma rate constants result in low translation loss. We also show that the time period of fluctuations in the environmental input can impose an additional criterion due to which central dogma rate constants exists in the relatively low integration time regime.

## Experimental system

To empirically evaluate information transfer in central dogma systems, we used recently reported single-cell measurements of inducible transcript and protein expression for the same gene [22, 23]. The experimental measurements were made using a system for Isopropyl *β*-D-1-thiogalactopyranoside(IPTG)-inducible expression of fluorescent protein in *E. coli*. For this system, IPTG serves as the environmental input which regulates the expression of enhanced yellow fluorescent protein (eYFP) (Fig. 1a). Single-cell expression of the *eyfp* transcript was measured using single-molecule RNA fluorescence *in situ* hybridization (FISH). Single-cell expression of the eYFP protein was measured using flow cytometry. Both methods were performed in parallel using cells harvested from the same culture, eliminating effects from culture-to-culture variation. These measurements of transcript and protein expression levels within the same system should capture the change in information as it transfers from the environment to the transcript expression level and then to the protein expression level.

**FIG. 1.**
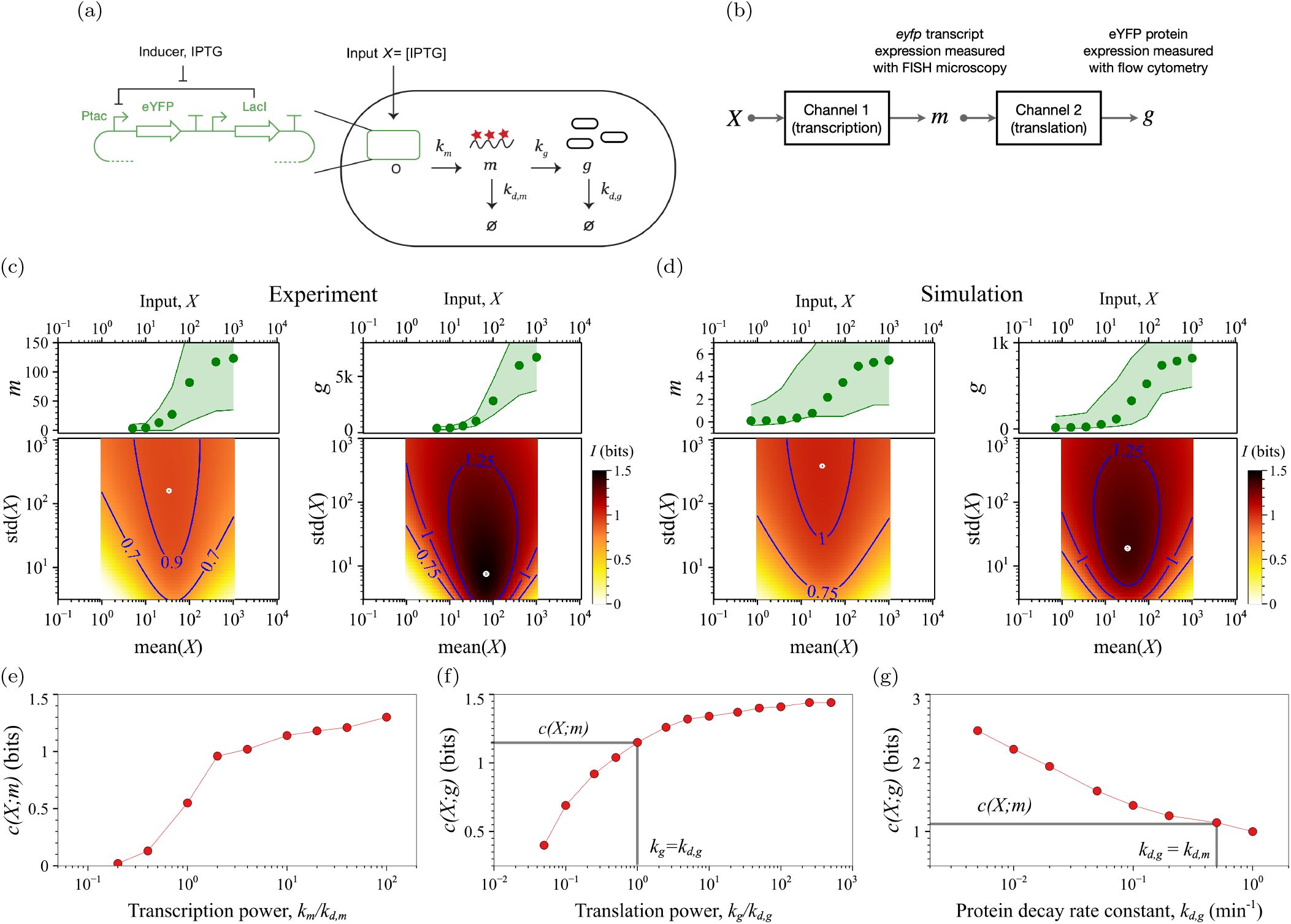
Information flow during transcription and translation. (a) Diagram of the experimental system where IPTG induces the expression of eYFP, for which we measure both the transcript and the protein expression levels. The central dogma rate constants consists of transcription (*k*_*m*_), translation (*k*_*g*_), and transcript and protein decays (*k*_*d,m*_, *k*_*d,g*_) [20]. (b) Sequential channel model of the central dogma process, which takes an input signal, *X*, and produces *eyfp* transcripts, *m*, as the output from the first channel and then eYFP proteins, *g*, as the output from the second channel. (c) Experimental result for the transcript-level mutual information (*I*(*X*; *m*), left), and the protein-level mutual information (*I*(*X*; *g*), right) using data from [22]. (d) Transcript-level (left) and protein-level (right) mutual information calculated from the simulation results, using the *lac operon*-based reaction network in [25]. In (c) and (d), the green dots in the top panels show the average transcript or protein expression levels, while the shaded region bounds the 5% to 95% percentiles; the 2D heat maps in the bottom panels show the mutual information values over the space of probability distributions of the input, *P* (*X*). The white dots in the heat map indicate the maximum mutual information. In (c) the transcript-level dose response curve is in the unit of RNA counts/cell and the protein-level dose response curves are in the units of molecules of equivalent fluorescein [43]. In (d) the dose-response curves are in the units of number of transcript and protein molecules per cell from Gillespie simulations. (e) Transcript-level channel capacity for varying transcription rate constant with fixed transcript decay rate constant, *k*_*d,m*_ = 0.5 min^−1^ (other rate constants the same as reported in [25]). (f) Protein-level channel capacity for varying translation rate constant with fixed protein decay rate constant, *k*_*d,g*_ = 0.2 min^−1^ (other rate constants the same as reported in [25]). As highlighted by the black lines, when translation power is 1, we find that the transcript and the protein-level channel capacities are approximately equal. (g) Protein-level channel capacity for varying protein decay rate constant with fixed translation power (*k*_*g*_*/k*_*d,g*_ = 10^3^) (other rate constants the same as reported in [25]). As highlighted by the black lines, when the transcript and the protein decay rates are the same, we find that the transcript and the protein-level channel capacities are approximately equal.

## Quantification of information transfer

To quantify the transfer of information in the experimental system, we consider transcription and translation as information channels, and we determine the mutual information between the environmental input and the transcript or the protein expression level (SI: 1). To understand information transfer through the central dogma processes, we compare the two quantities: mutual information between the environmental input and the transcript expression level (the transcript-level mutual information), and mutual information between the environmental input and the protein expression level (the protein-level mutual information). Mutual information is an entropic measure of the dependence between two random variables, in this case the input and either the transcript or the protein expression level [21, 24]. Mutual information associated with an information channel depends on both the transition matrix of the channel (*P*(*output*|*input*)) and the probability distribution of input signals. So, for a metric that characterizes the information channel alone, we consider the channel capacity, which is the maximum mutual information over all possible input distributions [12, 21].

Assuming that translation acts as a simple information channel that only preserves or degrades the information received from the transcription channel (Fig. 1b), we expected the protein-level channel capacity to be comparable to or lower than the transcript-level channel capacity. Surprisingly, we found the opposite: the experimentally observed protein-level channel capacity (*≈* 1.5 bits) is higher than the transcript-level channel capacity (*≈* 1.0 bits, Fig. 1c). Hence, there exists an apparent gain in information about the environmental input between the transcription and translation channels. Moreover, after evaluating both the transcript-level and protein-level mutual information for a large set of input distributions, we found that the protein expression level always contains higher information about the input compared to the transcript expression level (Fig. 1c, SI: 1A and 1C).

The observed information gain in the protein-level mutual information and channel capacity could be due to two factors other than the central dogma process itself: (1) the transcript expression measurement could be noisier than the protein expression measurement, and (2) there could be unknown biochemical reaction pathways that form an additional information channel parallel to transcription and translation and that transfers information directly from the input to protein expression, bypassing translation. To show that the gain in the protein-level information is a characteristic property of the central dogma rather than a result of measurement noise or unknown reaction pathways, we performed Gillespie (kinetic Monte Carlo) simulations of a biochemical reaction network that represents the experimental gene expression system [25] (SI: 1B). The simulated biochemical reaction network contains no unknown reaction pathways. More specifically, it only contains sequential transcription and translation processes and excludes any reaction pathway that could directly transfer information from the input to the protein expression level. Also, the simulation directly provides transcript and protein expression levels, excluding measurement noise as a possible factor. The Gillespie simulations were consistent with the experimental results: the protein-level mutual information is greater than the transcript-level mutual information for all input distributions considered (Fig. 1d and SI: 1C). We will show that the gain in the protein-level mutual information occurs due to time integration of the transcript expression level, and also depends on the translation power.

To explore how the central dogma rate constants affect the transfer of information from the input to the transcript and protein expression levels, we used additional Gillespie simulations to determine parametric trends in the transcript-level and protein-level channel capacities. We observed that the transcript-level channel capacity increases with increasing transcription power (the ratio of the transcription rate constant to the transcript decay rate constant, Fig. 1e), and that the protein-level channel capacity increases with increasing translation power (the ratio of the translation rate constant to the protein decay rate constant, Fig. 1f). These trends are similar to the property of simple information channels, *e*.*g*. Gaussian or Poisson channels, where channel capacity increases with channel power [21, 26, 27]. When the translation power is 1, then the protein-level channel capacity is close to the transcript-level channel capacity, and higher values of translation power appears to increase the protein-level channel capacity towards an asymptotic value (Fig. 1f). We will show that this asymptotic value depends on the ratio of the transcript decay rate constant to the protein decay rate constant. Interestingly, at fixed transcription and translation powers (*i*.*e*., fixed mean protein expression level), the protein-level channel capacity *decreases* with increasing protein decay rate constant (Fig. 1g). So, the transcript-level channel capacity increases with the transcription power, and the *increase* in the protein-level level channel capacity (*i*.*e*. the amount of information gained during translation) depends on the transcript and protein decay rate constants and the translation power.

## Channel model

To develop an information theoretic model that explains the information gain during translation, we start by considering a fundamental result in information theory: Information about the input can only be degraded as it transfers through each information channel. From this, it has been proven that if two information channels are connected in series, then the channel capacity of the combined channel must be less than the channel capacity of the first channel [21, 28]. However, that result is only true for “delayless processing”, in which the second channel only receives one symbol at a time from the first channel to produce a response (*i*.*e*., there is no accumulation of the first channel’s output by the second channel [29, 30]). In the context of the central dogma, the transfer of information from transcription to translation is only delayless if the response times for transcription and translation are equal. In general, however, those response times can be different.

To examine how the difference in the two response times can produce a gain in information during translation, we used a generic but sufficient model for transcription and translation that includes the four central dogma rate constants: transcription, transcript decay, translation, and protein decay (SI: 2). Within the generic model, the transcript expression is a stochastic process governed by the master equation,

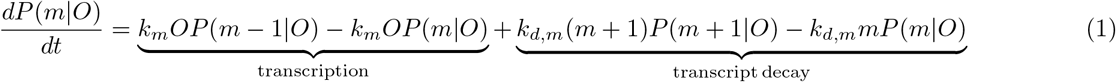

where *m* is the number of transcripts, *O* is the promoter state which is either on (*O* = 1) or off (*O* = 0), *k*_*m*_ is the transcription rate constant, and *k*_*d,m*_ is the transcript decay rate constant. Both transcription and transcript decay processes can increase or decrease the occupation probability of the transcript expression level, *P* (*m*|*O*). The protein expression is a stochastic process governed by the master equation,

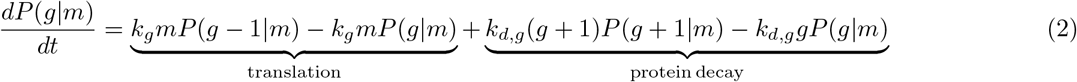

where *g* is the number of proteins, *k*_*g*_ is the translation rate constant and *k*_*d,g*_ is the protein decay rate constant, which effectively represents both active protein degradation as well as dilution due to cell division. From the deterministic ODEs that are obtained by ensemble-averaging the master equations (1) and (2) (SI: 2A and 2B), the response times for transcription and translation are 1*/k*_*d,m*_ and 1*/k*_*d,g*_, respectively. Hence, delayless processing in the central dogma requires *k*_*d,m*_ = *k*_*d,g*_. However, the transcript decay rate constant is generally faster than the protein decay rate constant, typically by a factor of 10 [20, 31–37]. Consequently, the protein response time is slower than the transcript response time, and the translation channel effectively receives and integrates multiple signals from the transcription channel.

## Maximum possible information gain during translation

To determine the amount of information lost due to stochasticity in translation, we first calculated the maximum possible information gain due to time integration during translation. From the ODE for the ensemble-averaged protein expression level we can show that the protein expression, *g*(*t*), is the convolution of the transcript trajectory, *m*(*t*), with a time integration kernel, 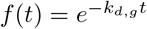 and multiplied by the translation rate constant *k*_*g*_ (SI: 2C and 2D). In the deterministic model of time integration during translation, *k*_*g*_ only scales the convolution output, without increasing the dispersion in the protein expression level, and therefore does not affect the protein-level channel capacity. So, the result of an ideal, noise-free time integration during translation is the hypothetical protein expression level: *g*_ideal_(*t*) *≡* (*f ∗ m*)(*t*). We define the maximum information transferred to this hypothetical protein expression level as the ideal channel capacity *c*_ideal_(*T*) *≡ c*(*X*; *g*_ideal_), as a function of the dimensionless integration time, *T ≡ k*_*d,m*_*/k*_*d,g*_. Furthermore, since the analytical solution to the transcript expression distribution, *P*(*m*|*X*), is known [6, 38], we can construct an analytical approximation of *g*_ideal_. The number of uncorrelated outputs of the transcription channel received by the translation channel within its response time is *T*. Therefore, we approximate 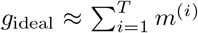, where each *m*^(*i*)^ are independent and identically distributed random variables with the distribution *P* (*m*|*X*). We found that the ideal channel capacity from the analytical approximation matches the value from numerical convolution of the transcript trajectory (SI: 2E).

We identified the combined effect of *k*_*m*_, *k*_*d,m*_, and *k*_*d,g*_ on *c*_ideal_(*T*) using multiple simulated data sets (Fig. 2a, SI: 3A). First we determined the transcript expression distribution as a function of *k*_*m*_ and *k*_*d,m*_, then we determined the ideal integration output distribution, *P*(*g*_ideal_|*X*), using the analytical approximation to compute *c*_ideal_(*T*) (Fig. 2a, SI: 2F and 3A). At *T* = 1 the ideal channel capacity is equal to the transcription channel capacity, and subsequent increase in the integration time *T* increases the maximum possible gain during translation. Stochasticity in translation reduces the protein-level channel capacity, but this reduction is against the ideal channel capacity and *not* against the transcript-level channel capacity.

**FIG. 2.**
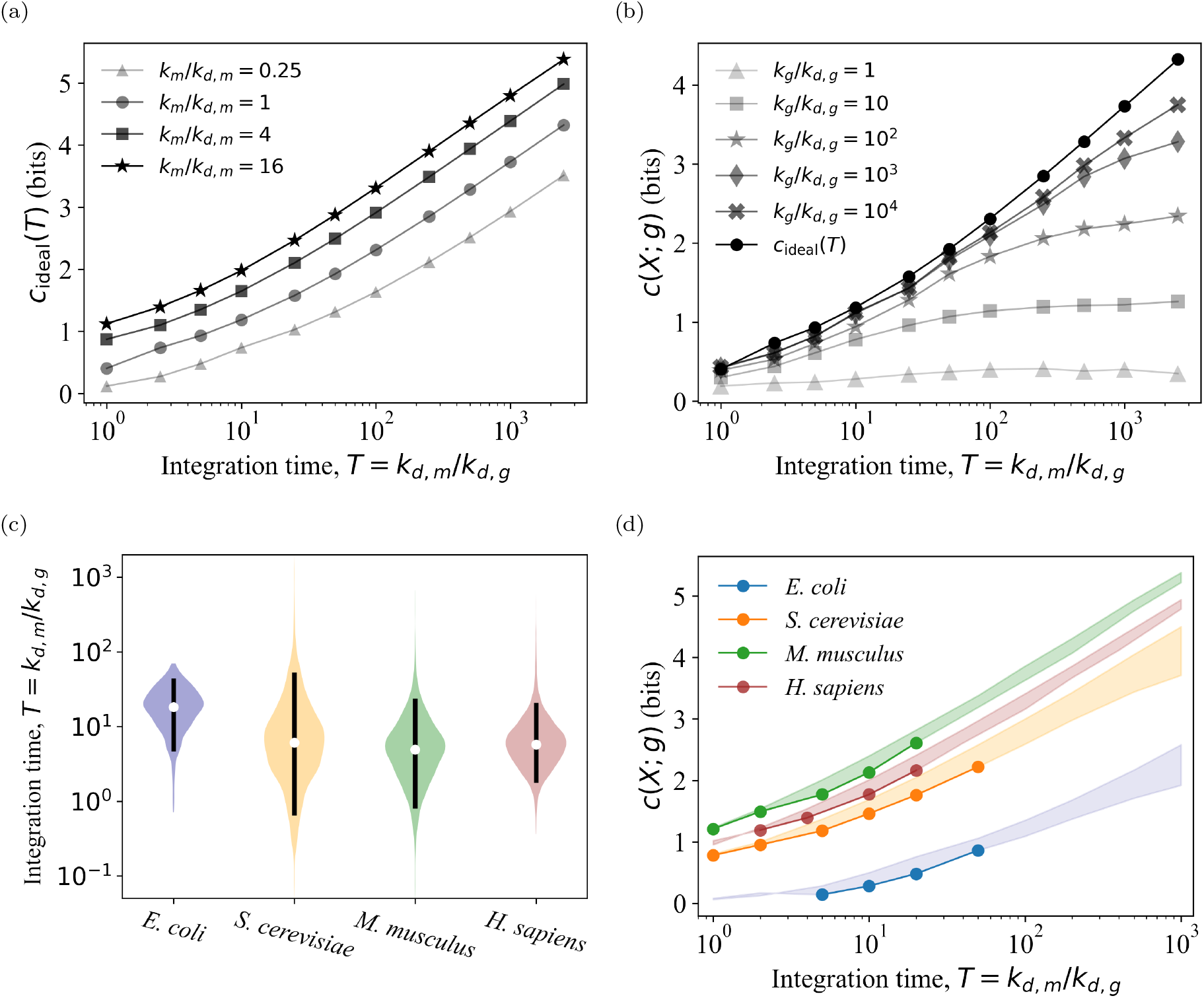
Information gain during translation. (a) Ideal information gain curves as a function of the dimensionless integration time, *T k*_*d,m*_*/k*_*d,g*_. At *T* = 1, the ideal channel capacity is a function of the transcription power, *k*_*m*_*/k*_*d,m*_. Results are shown for a fixed transcript decay rate constant *k*_*d,m*_ = 0.5 min^−1^. For *T >* 1, the ideal channel capacity, *c*_ideal_(*T*), increases, and is generally greater than the transcript-level channel capacity, *c*(*X*; *m*). (b) Protein-level information gain curves for different translation power values, *k*_*g*_*/k*_*d,g*_. For the results shown, *k*_*m*_ = *k*_*d,m*_ = 0.5 min^−1^, which results in relatively low transcript-level channel capacity, *c*(*X*; *m*) = 0.4 bits. The information gain due to time integration is determined by the translation channel power, *k*_*g*_*/k*_*d,g*_, and the dimensionless integration time. (c) Distributions of the dimensionless integration time for four species from data in published literature. The vertical black bars in each of the violin plots shows the 5% to 95% percentile range. The white dot in the middle of each black bar is the median. (d) Ideal and protein-level information gain curves for the four species. The solid lines with dots shows the protein-level information gain curve for each species, *c*(*X*; *g*), approximately covering the 5% to 95% percentile range of their respective integration time distributions. The shaded regions highlight the translation loss, which is the difference between the maximum possible information transfer and the actual information transfer for the same integration time, *c*_ideal_(*T*) − *c*(*X*; *g*). The plot shows the translation loss both within and beyond the naturally occurring range of integration times.

## Information lost due to stochasticity in translation

To determine the loss in information after time integration, we estimated the true protein-level channel capacity from the distributions *P*(*g*|*X*) obtained using stochastic simulations of equations (1) and (2). We computed the protein-level information gain curves, *c*(*X*; *g*) vs. *T*, for five values of the translation power, *k*_*g*_*/k*_*d,g*_, which determines the mean protein expression level (SI: 3A). At low translation power, there is relatively more noise in the translation output, and the protein-level information gain curve becomes significantly lower than the ideal information gain curve, *c*_ideal_(*T*) vs. *T* (Fig. 2b). As the translation power increases, the protein-level channel capacity asymptotically approaches the ideal channel capacity, *c*_ideal_(*T*).

We observed two generic features in the information gain curves. First, the protein-level channel capacity, *c*(*X*; *g*), increases monotonically with integration time but has a plateau at higher integration times. This plateau is most prominent for translation power *k*_*g*_*/k*_*d,g*_ ≤ 100 (Fig. 2b). Increasing the translation power shifts the onset of the plateau region to higher integration times. Second, the translation loss, *i*.*e. c*_ideal_(*T*) − *c*(*X*; *g*), is generally small for low integration times (before the plateau region). But in the plateau region, the translation loss increases dramatically with increasing integration time. Hence, although higher integration time can produce higher information gain it also causes a higher translation loss.

## Information gain in naturally evolved systems

To estimate the information gain due to time integration and the translation loss for naturally evolved systems, we performed stochastic simulations of the central dogma process using typical rate constants for four species: *E. coli, S. cerevisiae, M. musculus*, and *H. sapiens*. For each of the species, we used values for the central dogma rate constants from published data (SI: 3B) [20, 31–37]. We used those values to determine the distribution of the dimensionless integration time for each of the species (Fig. 2c), and then to compute the ideal information gain curves, *c*_ideal_(*T*). Similarly, we used the distributions resulting from stochastic simulations of transcript and protein expression levels to compute the protein-level information gain curves (Fig. 2d). The median value of the dimensionless integration time for *E. coli* is 20 (5% to 95% percentile range: 5 to 44). For eukaryotic species, the median dimensionless integration time is lower: 6 (1 to 53) for *S. cerevisiae*, 5 (1 to 24) for *M. musculus*, and 6 (2 to 21) for *H. sapiens* (Fig. 2c, SI: 3B). We do not have similarly extensive data for other bacterial species, but the typical integration time for *B. subtilis* also appears to around 20 [39, 40]. So, prokaryotes may generally have higher dimensionless integration times than eukaryotes. Based on the gene ontology enrichment analysis for the *M. musculus* data (using paired protein and transcript decay rates reported in the same study) [35]: genes with relatively high dimensionless integration times are associated with dephosphorylation and RNA processing. Genes with relatively low dimensionless integration times are associated with defense response, homeostasis, and proteolysis — processes that may require faster response times.

For all four species, within the typical range of integration times (the 5% to 95% percentile range of *T*), the translation loss is less than 0.5 bits (Fig. 2d), which means the translation power, or the average protein expression level, is nearly adequate to transfer the maximum amount of information enabled by the time integration. This suggests that low translation loss could be an evolutionary selection criterion for the central dogma rate constants. From our simulation results (Fig. 2b), some combinations of central dogma rate constants can result in high translation loss (*>* 1 bit), but we do not observe such rate constant combinations among the values reported in the literature. As a corollary to the observation of low translation loss, the naturally occurring central dogma rate constants do not achieve the plateauing or maximum information gain possible for the given translation power. So, maximizing the information gain for a fixed average protein expression level is probably not a selection criterion. Moreover, although the translation loss appears to be consistently low across species, the observed integration times do not span the full range of low translation loss: they stop well below the onset of the plateau region (Fig. 2d). Hence, although low translation loss might be an evolutionary selection criterion, it does not appear to be the only (or even the dominant) criterion.

To understand why naturally evolved systems do not have dimensionless integration times greater than approximately 100, we considered the time scale of fluctuations in the environmental input. High integration times correspond to slower translation response times, and central dogma systems can only operate at channel capacity if the environmental fluctuations are slower than the translation response time (SI: 3C). When the input fluctuation period is less than or comparable to the translation response time, then the translation channel output is correlated with the previous outputs. These correlations decrease the channel capacity in a way that is analogous to the reduced capacity of slow-fading information channels [41, 42]. Using the ideal information gain model, we found that environmental fluctuations have to occur with a period roughly 10 times longer than the integration time for the protein-level mutual information *I*(*X*; *g*) to be close to capacity. Thus, we speculate that the four species we examined and other naturally evolved biological systems have central dogma rate constants that use time-integration for information gain but remain within the regime of negligible translation loss and relatively fast translation response times.

## Conclusion

We developed an information gain model for central dogma systems in biology that describes the flow of information from the environment to the transcript expression level and then to the protein expression level. This information gain model explains the mutual information values found in single-cell measurements and stochastic simulations. Results of the information gain model with central dogma rate constants typical for four different species indicate that information gain from time integration during translation is a general feature that results in a gain of information at the protein expression level. Furthermore, for all four species, the central dogma rate constants are typically found only in a regime where the information loss due to stochasticity in translation is relatively low. This suggests that time integration with low translation loss and avoiding very slow response times may be selection criteria for naturally evolved central dogma systems.

Our findings regarding the distributions of integration times, the protein-level information gain, and the translation loss in the four species (Fig. 2) raise questions for future research in biochemical signaling. For example, the literature results for decay rates suggest that the typical dimensionless integration time is lower in eukaryotes than prokaryotes. Paired transcript and protein decay rate data for additional species would help confirm whether this trend is universal. Also, since the translation loss is small for the four species, the ideal channel capacity described here can provide a fast estimate of the protein-level channel capacity that does not require protein expression data. This could be a useful approximation for large surveys of biological information transfer.

Beyond transforming the central dogma process from a set of biochemical reactions to an information acquiring and integrating system, these insights are also relevant for engineering synthetic biological information processing systems. Synthetic biology enables the construction of gene expression systems with a wide range of central dogma rate constants. This work presents new quantitative criteria, the translation loss and the response time relative to the time scale of environmental fluctuations, as essential factors to consider for the design of synthetic central dogma systems.

## Code availability

Source code to compute the ideal and the protein-level information gain curves, and the transcript and protein decay rates data are available at github.com/sarkar-s/InCens.

## Acknowledgements

We would like to especially thank David Ross for constructive discussions during the research and preparation of this manuscript. We would also like to thank Samuel Schaffter, Peter Tonner, Elizabeth Strychalski, and Charles Camp for thoughtful feedback on this manuscript.

