## Supplementary Information for "Nearly maximal information gain due to time integration in central dogma reactions"

### CONTENTS

SI: 1. Computation of mutual information and channel capacity 1
A. Transcript and protein-level mutual information landscapes for biological replicates 2
B. Simulated inducible gene expression system 2
C. Difference in the transcript and protein-level mutual information 3

SI: 2. Generic stochastic model of transcription and translation 3
A. Master equations 4
B. Governing equations for mean transcript and protein expression levels 4
C. Translation output for a time-dependent transcript expression 5
D. Deterministic integration approximation for the translation output 6
E. Validation of the analytical approximation for ideal channel capacity 7
F. Effect of number of input levels on the estimate of $c_{\text{ideal}}(T)$ 8

SI: 3. Stochastic simulations of central dogma master equations 8
A. Parameters for Fig. 2(a) and 2(b) 8
B. Parameters for the four species, for Fig. 2(c) and 2(d) 9
C. Effect of fluctuation time period on information transfer 11

References 12

### SI: 1. COMPUTATION OF MUTUAL INFORMATION AND CHANNEL CAPACITY

To assess the transfer of information from the environmental input to either the transcript or the protein expression levels we computed the mutual information in bits as

$$I(X; g) = \sum_X P(X) \sum_g P(g|X) \log_2 \frac{P(g|X)}{P(g)} \tag{1}$$

For the transcript level mutual information, we replace the protein expression level $g$ with the transcript expression level $m$ in Eq. (1). The conditional distributions, $P(g|X)$ or $P(m|X)$, in Eq. (1), are empirical distributions of the transcript or protein expression levels for fixed values of the environmental input, either from experiments or from stochastic simulations. The set of transcript or protein expression distributions, $P(m|X)$ or $P(g|X)$, for a set of values of the input, $X$, mathematically defines the information channel for which we compute the mutual information. As seen in Eq. (1), mutual information also depends on the input distribution, $P(X)$. The maximum possible mutual information through an information channel for all possible input distributions is the channel capacity, which is calculated by maximizing the mutual information with respect to the input distribution:

$$c(X; g) = \max_{P(X)} I(X; g) \tag{2}$$

Throughout this work we computed the channel capacity using the well-established Blahut-Arimoto algorithm, as described in [1–4], and then corrected for finite-sampling bias as described in [2, 3]. The mutual information landscapes shown in Fig. 1(c) and 1(d) were computed from the transcript and the protein expression distributions, $P(m|X)$ and $P(g|X)$, respectively, using Sparse Estimation of Mutual Information Landscapes (SEMIL) as described in [3].

*

#### A. Transcript and protein-level mutual information landscapes for biological replicates

To check the reproducibility of the transcript and the protein-level mutual information values we computed the mutual information landscapes with data from 3 biological replicates of the inducible gene expression system [5]. As mentioned in the main text, the transcript-level expression was measured using microscopy of FISH probes, and the protein-level expression was measured using flow cytometry [6]. The transcript and the protein-level mutual information landscapes for the 3 replicates are shown in Fig. SI: 1. The landscapes show the mutual information values across a space of input distributions, $P(X)$, where the distributions are identified using the mean and the standard deviation of $P(X)$.

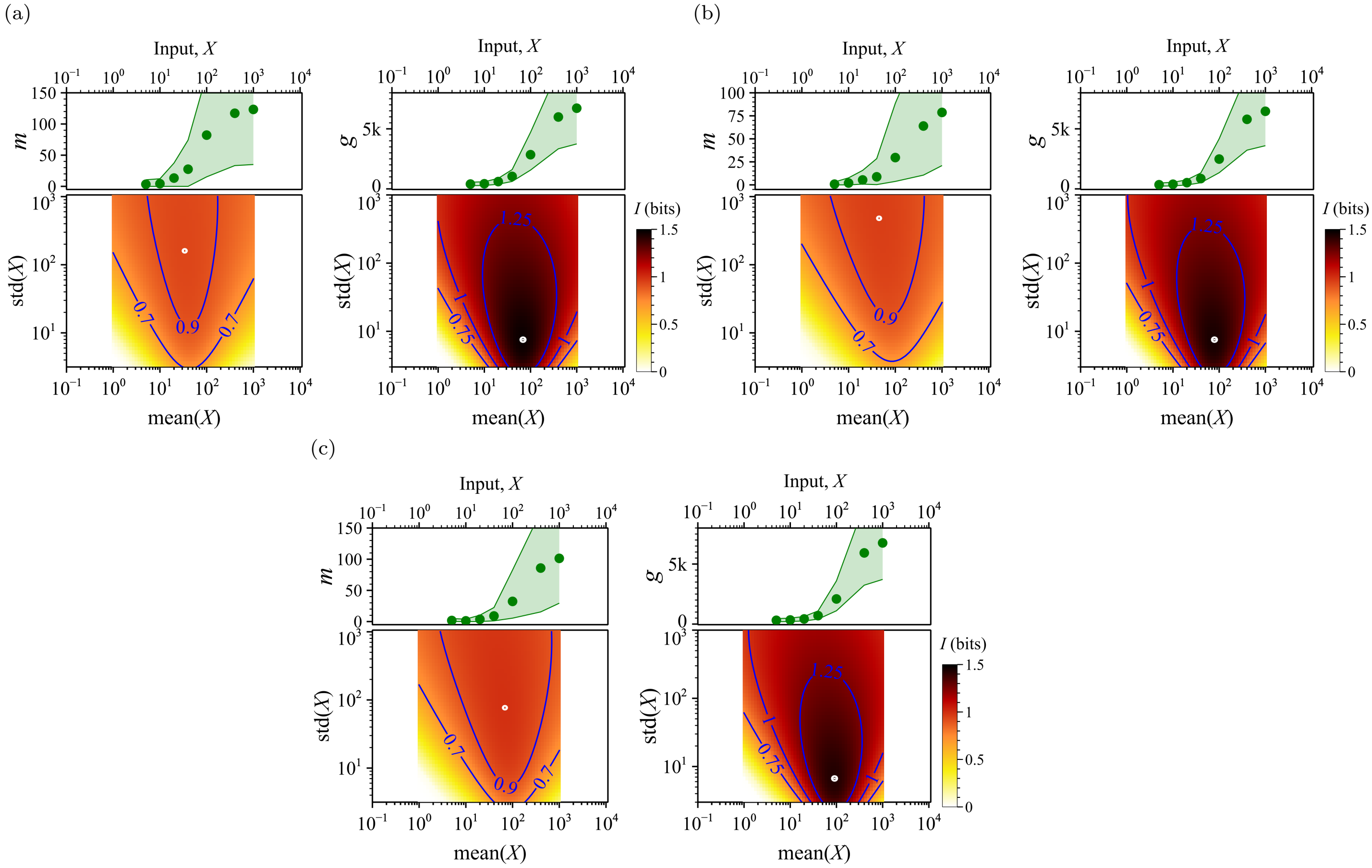


FIG. SI: 1. Transcript and protein-level mutual information landscapes for the three biological replicates measured experimentally: (a) replicate 1, (b) replicate 2, and (c) replicate 3. The transcript and the protein-level dose response curves (in green) and the mutual information landscapes (heat map) have the same annotation as in Fig. 1(c).

#### B. Simulated inducible gene expression system

To check that the central dogma reactions, consisting only of sequential transcription and translation, can cause the gain in the protein-level mutual information, we computed the mutual information landscapes for a simulated reaction network that is analogous to our experimental system. This reaction network was developed from a previously-published model of the *lac operon* [3, 7], consisting of the same set of reactions involving the input (IPTG) and the operator as present in the *lac operon* but without the positive feedback due to the *lacY* gene, as our experimental system does not contain feedback. We used Gillespie simulations to compute the transcript and the protein expression levels for this model reaction network. We used exactly the same set of reactions and rate constants shown in the SI of [3] in Tables 4 and 5, respectively. We subsequently used the transcript and the protein expression data to compute the mutual information landscapes as described in [3]. The transcript and the protein-level mutual information landscapes for the simulated inducible gene expression system are shown in Fig. 1(d) of the main text.

#### C. Difference in the transcript and protein-level mutual information

To show that the gain in the protein-level mutual information exists for all input distributions, we present the difference between the protein-level and transcript-level mutual information landscapes in Fig. SI: 2. The protein-level mutual information is higher than the transcript-level mutual information for all the input distributions in the landscape, so the gain in protein-level mutual information is not confined to the input probability distribution that causes maximum information transfer, but is most likely true for all input distributions.

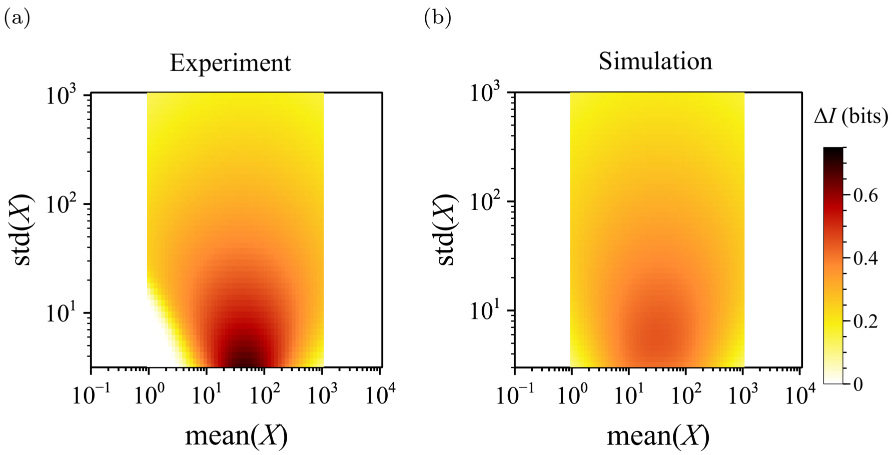


FIG. SI: 2. Difference between the protein-level and the transcript-level mutual information, $\Delta I = I(X; g) - I(X; m)$, from both the experimental and the simulated data. The difference in mutual information values are shown over the space of probability distributions of the input. Input distributions, $P(X)$, are identified in the landscape using the mean and the standard deviation as a two dimensional coordinate. (a) Gain in the protein-level mutual information, $I(X; g)$, compared to the transcript-level mutual information, $I(X; m)$, for the experimental inducible gene expression system shown in Fig. 1(a) of the manuscript. (b) Gain in the protein-level mutual information, $I(X; g)$, compared to the transcript-level mutual information, $I(X; m)$, for the simulated inducible gene expression system [3].

### SI: 2. GENERIC STOCHASTIC MODEL OF TRANSCRIPTION AND TRANSLATION

To generate the stochastic transcript and protein expression data for a generic central dogma system, we considered that the information transfer from the environment to the protein expression level occurs through the following three processes.

1. The operator state switching between active ($O = 1$) and inactive ($O = 0$).

$$(O = 1) \underset{k_{OFF}}{\overset{k_{ON}}{\rightleftharpoons}} (O = 0) \tag{3}$$

2. Transcription and transcript decay.

$$\begin{aligned} (O = 1) &\xrightarrow{k_m} (O = 1) + m \\ m &\xrightarrow{k_{d,m}} \emptyset \end{aligned} \tag{4}$$

3. Translation and protein decay.

$$\begin{aligned} m &\xrightarrow{k_g} m + g \\ g &\xrightarrow{k_{d,g}} \emptyset \end{aligned} \tag{5}$$

The set of processes, (3)-(5), is a sufficient but minimal model of the central dogma system. The interaction of the environmental input with the operator is condensed into the rate constants, $k_{ON}$ and $k_{OFF}$. In this work, we consider the following form for the $k_{ON}$ and $k_{OFF}$ rate constants

$$\begin{aligned} k_{ON} &= \alpha((1-l)X + l) \\ k_{OFF} &= \alpha(1-l)(1-X) \end{aligned} \tag{6}$$

where $0 \leq X \leq 1$ is the environmental input value. $l$ is the leakiness, or $l = k_{ON}/(k_{ON} + k_{OFF})$ when $X = 0$, which determines the leaky transcription in the absence of the environmental input. $\alpha$ controls the rate or the frequency of switching between active and inactive operator states, as $k_{ON} + k_{OFF} = \alpha$ independent of the input value $X$. The transcript and the protein-level channel capacity depends only on the expression distributions, $P(m|X)$ and $P(g|X)$, respectively. The shape of mean dose-response curves, $\langle m \rangle$-vs-$X$ or $\langle g \rangle$-vs-$X$, does not change the channel capacity, as long as the distributions $P(m|X)$ and $P(g|X)$ remain unchanged.

#### A. Master equations

The master equation for the active state of the operator is

$$\frac{dP(O=1)}{dt} = k_{ON}P(O=0) - k_{OFF}P(O=1) \tag{7}$$

Since the operator state can be either active or inactive at a time, the master equation for the inactive state of the operator is similar to Eq. (7), with the right hand side multiplied by -1, *i.e.*,

$$\frac{dP(O=0)}{dt} = k_{OFF}P(O=1) - k_{ON}P(O=0) \tag{8}$$

The master equation for the transcript expression level (or transcript copy number), $m$, is

$$\frac{dP(m|O)}{dt} = k_m O P(m-1|O) + k_{d,m}(m+1)P(m+1|O) - k_m O P(m|O) - k_{d,m} m P(m|O) \tag{9}$$

The master equation for the protein expression level (or protein copy number), $g$, for a fixed valued of transcript expression, $m$, is

$$\frac{dP(g|m)}{dt} = k_g m P(g-1|m) + k_{d,g}(g+1)P(g+1|m) - k_g m P(g|m) - k_{d,g} g P(g|m) \tag{10}$$

Both equations (9) and (10) are one-step master equations.

#### B. Governing equations for mean transcript and protein expression levels

The governing equation for the ensemble-averaged mean transcript expression level, $\langle m|O \rangle$, for a fixed operator state $O$ is obtained by multiplying the transcription master equation (9) with the transcript expression and summing over all possible values as

$$\begin{aligned} \frac{d\langle m|O \rangle}{dt} = \frac{d\sum_{m=0}^{\infty} mP(m|O)}{dt} =& k_m O \sum_{m=0}^{\infty} mP(m-1|O) + k_{d,m} \sum_{m=0}^{\infty} m(m+1)P(m+1|O) \\ &- k_m O \sum_{m=0}^{\infty} mP(m|O) - k_{d,m} \sum_{m=0}^{\infty} m^2 P(m|O) \end{aligned} \tag{11}$$

$$
\begin{aligned}
\frac{d\langle m|O\rangle}{dt} &= k_m O \sum_{m=0}^{\infty} mP(m-1|O) + k_{d,m} \sum_{m=0}^{\infty} m(m+1)P(m+1|O) - k_m O\langle m|O\rangle - k_{d,m}\langle m^2|O\rangle \\
&= k_m O \sum_{m=0}^{\infty} (m+1)P(m|O) + k_{d,m} \sum_{m=0}^{\infty} m(m-1)P(m|O) - k_m O\langle m|O\rangle - k_{d,m}\langle m^2|O\rangle \\
&= k_m O - k_{d,m}\langle m|O\rangle
\end{aligned} \tag{12}
$$

The solution to Eq. (12) is

$$\langle m|O\rangle(t) = \frac{k_m O}{k_{d,m}}(1 - e^{-k_{d,m}t}) \tag{13}$$

which has a relaxation time constant $\frac{1}{k_{d,m}}$.

The governing equation for the ensemble-averaged mean protein expression level, $\langle g|m\rangle$ for a fixed transcript expression level is obtained from the translation master equation as

$$
\begin{aligned}
\frac{d\langle g|m\rangle}{dt} = \frac{d\sum_{g=0}^{\infty} gP(g|m)}{dt} =& k_g m \sum_{g=0}^{\infty} gP(g-1|m) + k_{d,g} \sum_{g=0}^{\infty} g(g+1)P(g+1|m) \\
& - k_g m \sum_{g=0}^{\infty} gP(g|m) - k_{d,g} \sum_{g=0}^{\infty} g^2 P(g|m)
\end{aligned} \tag{14}
$$

$$
\begin{aligned}
\frac{d\langle g|m\rangle}{dt} &= k_g m \sum_{g=0}^{\infty} gP(g-1|m) + k_{d,g} \sum_{g=0}^{\infty} g(g+1)P(g+1|m) - k_g m\langle g|m\rangle - k_{d,g}\langle g^2|m\rangle \\
&= k_g m \sum_{g=0}^{\infty} (g+1)P(g|m) + k_{d,g} \sum_{g=0}^{\infty} g(g-1)P(g|m) - k_g m\langle g|m\rangle - k_{d,g}\langle g^2|m\rangle \\
&= k_g m - k_{d,g}\langle g|m\rangle
\end{aligned} \tag{15}
$$

The solution to Eq. (15) is

$$\langle g|m\rangle(t) = \frac{k_g m}{k_{d,g}}(1 - e^{-k_{d,g}t}) \tag{16}$$

which has a relaxation time constant $\frac{1}{k_{d,g}}$.

#### C. Translation output for a time-dependent transcript expression

Eq. (16) is the average protein expression level for a fixed transcript expression level. By ensemble-averaging the Eq. (15) with the transcript expression distribution $P(m)$, we obtain the familiar deterministic ODE for translation as

$$\frac{d\langle g\rangle}{dt} = k_g\langle m\rangle - k_{d,g}\langle g\rangle \tag{17}$$

where both $\langle g\rangle$ and $\langle m\rangle$ are time-dependent. We can obtain the solution for a time-dependent average transcript expression level $\langle m\rangle(t)$ using Laplace transform. Let the Laplace transform of $\langle m\rangle(t)$ be $M(s)$ and of $\langle g\rangle(t)$ be $G(s)$, then the Laplace transform of Eq. (17) is

$$sG(s) = k_g M(s) - k_{d,g}G(s) \tag{18}$$

with the initial condition $\langle g\rangle(0^-) = 0$. Therefore,

$$G(s) = M(s)\frac{k_g}{s + k_{d,g}}$$
$$\langle g\rangle(t) = k_g \int_0^t \langle m\rangle(\tau) e^{-k_{d,g}(t-\tau)}\, d\tau \tag{19}$$

#### D. Deterministic integration approximation for the translation output

Eq. (19) is the deterministic model of translation, which neglects the stochasticity associated with the translation and the protein decay processes. We use the convolution operator from Eq. (19) to define the output of the deterministic integration of a *stochastic* transcript expression trajectory as

$$g_{\text{ideal}}(t) = (f * m)(t) = \int_0^t m(\tau) e^{-k_{d,g}(t-\tau)}\, d\tau \tag{20}$$

The convolution kernel $f$ accounts for the time integration of the stochastic transcript expression that occurs due to the response time of the translation process, $1/k_{d,g}$. We omit the translation rate constant $k_g$ from the operator in Eq. (19), because it only scales the output of the convolution without introducing more stochasticity to the output, which is necessary to have any impact on the information transfer. Example of the convolution output, which we have named the *ideal integration output*, as a function of the ratio $k_{d,m}/k_{d,g}$ is shown in Fig. SI: 3.

Using Eq. (20) we transform $m(t)$ to $g_{\text{ideal}}(t)$. From a transcript trajectory we obtain the transcript expression distribution, $P(m|X)$, and from the ideal integration output trajectory we obtain the distribution, $P(g_{\text{ideal}}|X)$. The channel capacity of the integrated output $c(X; g_{\text{ideal}})$ is a function of, $k_{d,g}$, or more specifically of the ratio $k_{d,m}/k_{d,g}$. $c(X; g_{\text{ideal}})$ as a function of the integration time $T = k_{d,m}/k_{d,g}$ is the information gain due to deterministic time integration during the translation process.

We constructed an analytical approximation of the ideal integration output of the transcript expression. We approximated that after every interval of the transcription response time, $1/k_{d,m}$, the transcript expression is represented using independent and identically distributed random variables, all with the distribution $P(m|X)$. Since the total number of intervals of $1/k_{d,m}$ during the translation response time is $T = k_{d,m}/k_{d,g}$, we define the ideal integration output as

$$g_{\text{ideal}}^{(\text{analyt.})} = \sum_{i=1}^{T} m^{(i)} \tag{21}$$

On the basis of Eq. (21), if the probability generating function of the transcript expression distribution is, $G(z; m)$, then the probability generating function of $g_{\text{ideal}}^{(\text{analyt.})}$ is $G(z; m)^T$. The transcript expression has either a negative binomial distribution (NB) or a Poisson distribution (Pois) depending on the Fano factor $\frac{\sigma^2(m)}{\langle m\rangle}$ [8]. If $m \sim \text{NB}(r, p)$, then $g_{\text{ideal}}^{(\text{analyt.})} \sim \text{NB}(rT, p)$, where $r$ is the number of failures and $p$ is the failure probability of the transcript expression distribution. Similarly, if $m \sim \text{Pois}(\lambda)$, then $g_{\text{ideal}}^{(\text{analyt.})} \sim \text{Pois}(\lambda T)$, where $\lambda$ is average transcript expression level. Although, $T$ is an integer in Eq. (21), we extend the use to real values while determining the negative binomial or Poisson distribution of the integrated output. The parameters for the transcript expression were calculated from $k_{ON}, k_{OFF}, k_m$, and $k_{d,m}$ using the method described in [8], with

$$\langle m\rangle = \frac{k_{ON}}{k_{ON} + k_{OFF}} \frac{k_m}{k_{d,m}}$$
$$b = \frac{\sigma^2}{\langle m\rangle} = 1 + \frac{k_{d,m} k_{OFF} \langle m\rangle}{k_{ON}(k_{ON} + k_{OFF} + k_{d,m})} \tag{22}$$

where $k_{ON}$ and $k_{OFF}$ were calculated for each $X$ from Eq. (6). When $b > 1$, then the transcript expression distribution is $\text{NB}(r, p)$, with $p = (b-1)/b$, and $r = \langle m\rangle/(b-1)$. When $b = 1$, then transcript expression distribution is $\text{Pois}(\lambda)$ with $\lambda = \langle m\rangle$.

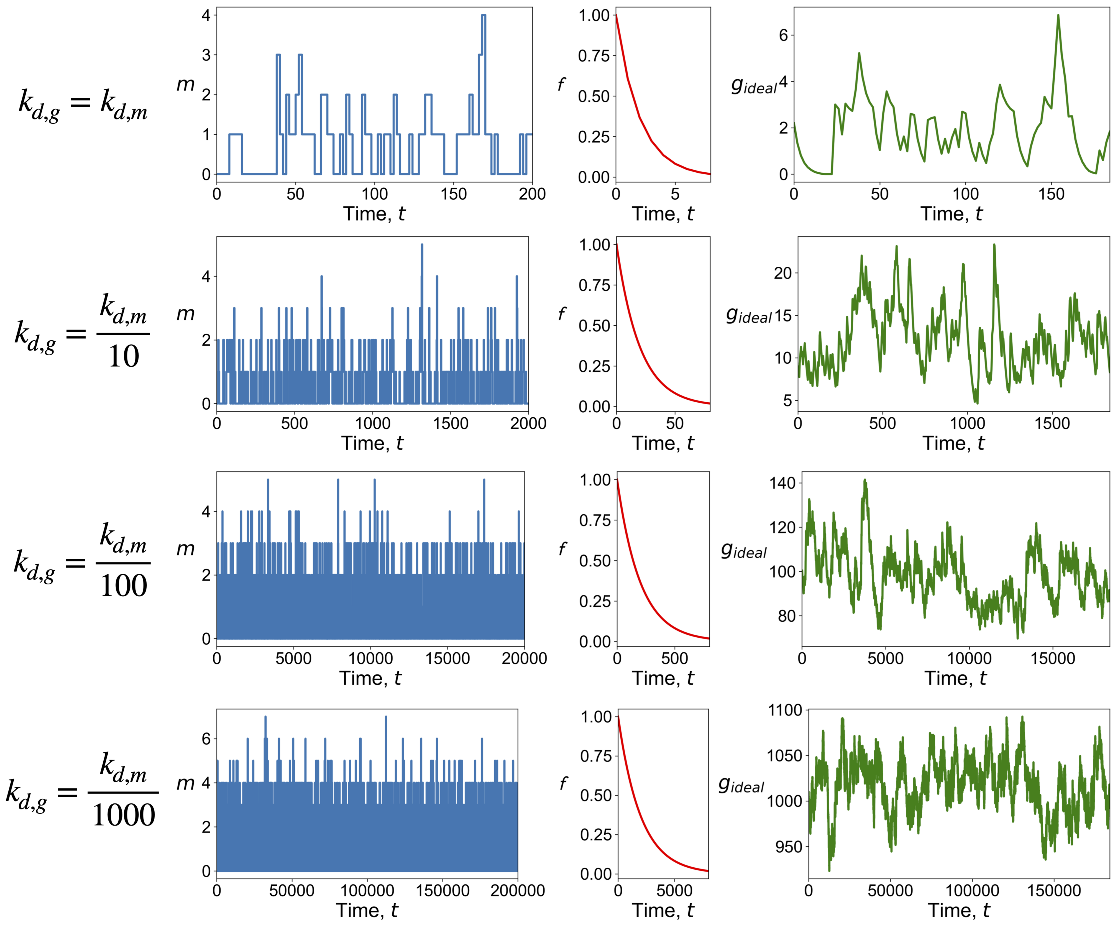


FIG. SI: 3. Integration of the stochastic time-dependent transcript expression level as a function of the protein decay rate constant $k_{d,g}$. The transcript decay rate constant is $k_{d,m} = 0.5\,\mathrm{min}^{-1}$ for all the examples, and the protein decay rate constant, $k_{d,g}$, is a decreasing fraction of the $k_{d,m}$ for each row of the figure. The stochastic transcript expression, $m(t)$, was captured for a duration of the $100/k_{d,g}$, sampled at interval of $1/k_{d,m}$. The convolution kernel function $f$ has the domain $[0, 4/k_{d,g}]$, with values at an interval of $1/k_{d,m}$. The output of the convolution $g_{\mathrm{ideal}}(t)$ is shown in green. The average $g_{\mathrm{ideal}}$ value increases as a function of increasing response time for translation, $1/k_{d,g}$, which is due to the increasing integration of the transcript expression level.

#### E. Validation of the analytical approximation for ideal channel capacity

To validate that the approximation, $g_{\mathrm{ideal}}^{(\mathrm{analyt.})}$, accurately estimates the ideal channel capacity, $c(X; g_{\mathrm{ideal}})$, we simulated a central dogma system with parameters, $k_m = k_{d,m} = 0.5\,\mathrm{min}^{-1}$, leakiness $l = 0.01$, frequency parameter $\alpha = 1.0\,\mathrm{min}^{-1}$, and with the input values $X = \{0.0, 0.1, 0.2, 0.3, 0.4, 0.5, 0.6, 0.7, 0.8, 0.9, 1\}$. We selected a set of protein decay rate constants, $k_{d,g} = \{0.5, 0.2, 0.1, 0.05, 0.02, 0.01, 0.005, 0.002, 0.001, 0.0005, 0.0002\}\mathrm{min}^{-1}$, which covers the range of integration times as $T = \{1, 2.5, 5, 10, 25, 50, 100, 250, 500, 1000, 2500\}$. For each $X$ we simulated the central dogma system up to transcription, Eq. (7)-(9), to obtain a stochastic transcript expression trajectory, $m(t)$. Then we obtained the time-integrated trajectory as $g_{\mathrm{ideal}}(t)$ using (20), which was then used to compute the channel capacity $c(X; g_{\mathrm{ideal}})$. At the same time we determined $P(m|X)$ using $k_{ON}, k_{OFF}, k_m$, and $k_{d,m}$ as described in [8]. We then computed the probability distribution of $g_{\mathrm{ideal}}^{(\mathrm{analyt.})}$ using (21) and subsequently determined $c(X; g_{\mathrm{ideal}}^{(\mathrm{analyt.})})$.

We performed two numerical case studies to check if $c(X; g_{\mathrm{ideal}}) = c(X; g_{\mathrm{ideal}}^{(\mathrm{analyt.})})$. In the first case, we determined the transcript stochastic trajectory for $10^4, 10^5$, and $10^6$ samples at an interval of $1/k_{d,m}$, and convoluted them with the kernel $f(t)$ which was represented in the time domain $[0, 4/k_{d,g}]$ with values at interval of $1/k_{d,m}$. We found that $c(X; g_{\mathrm{ideal}}) = c(X; g_{\mathrm{ideal}}^{(\mathrm{analyt.})})$ within 0.1 bits for all the 3 sizes of trajectory data across the entire range of integration time Fig. SI: 4a. In the second case, we chose the transcript trajectories with $10^5$ samples and convoluted with 4 representations of $f(t)$, with increasing time domains $[0, 1/k_{d,g}]$, $[0, 2/k_{d,g}]$, $[0, 4/k_{d,g}]$, and $[0, 8/k_{d,g}]$. We found that when the time domain is $[0, 4/k_{d,g}]$ or larger, then the ideal channel capacity from the numerical convolution matches with the analytical approximation (within 0.1 bits), Fig. SI: 4b. Based on the results in Fig. SI: 4, we conclude that for a sufficiently large transcript expression trajectory and convolution kernel domain $c(X; g_{\text{ideal}}) = c(X; g_{\text{ideal}}^{(\text{analyt.})})$.

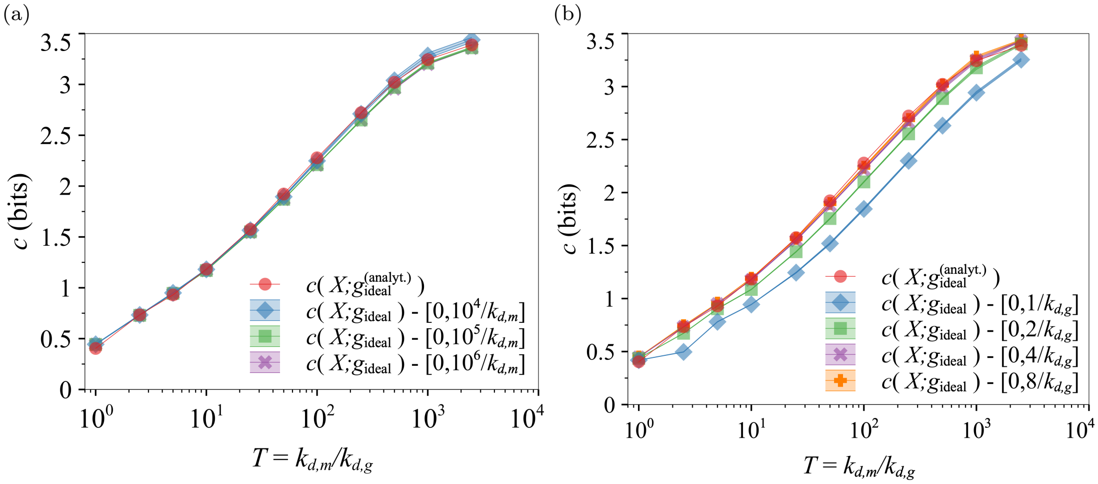


FIG. SI: 4. Estimate of the information gain due to integration using the numerical and analytical methods as described in SI: 2 D. (a) Effect of sample size or the length of the transcript expression trajectory $m(t)$ on the estimate of the information gain curve $c(X; g_{\text{ideal}})$-vs-$T$. The 95% confidence interval for each channel capacities $c(X; g_{\text{ideal}})$ was obtained from 20 replicates. (b) Effect of the time domain size for the kernel function $f(t)$ on the estimate of the information gain curve $c(X; g_{\text{ideal}})$-vs-$T$. The 95% confidence interval for each channel capacities $c(X; g_{\text{ideal}})$ was obtained from 10 replicates. In both (a) and (b) the red solid line with dots show the ideal information gain using the analytical approximation to the ideal integration output, Eq. (21).

#### F. Effect of number of input levels on the estimate of $c_{\text{ideal}}(T)$

The estimate of the ideal channel capacity is bounded from above by $\log_2 z$, where $z$ is the number of input values $X$ for which we have the distributions $P(g_{\text{ideal}}|X)$, either from numerical convolution or analytical approximation. Since we used 11 values of $X$ for Fig. SI: 4, all the information gain curves peak at $\log_2 11 \approx 3.5$ bits. However, the correct value of $c_{\text{ideal}}(T)$ is not bounded by the number of input values. To remove the underestimation of $c_{\text{ideal}}(T)$ due to the number of input values, we systemically increased the number of input levels as $z \in \{4, 8, 16, 32, 64, 128\}$ for $X \in [0, 1]$, which increases the entropy of the input as $H(X) \in \{2, 3, 4, 5, 6, 7\}$ bits, respectively. For each set of input values, we computed $c_{\text{ideal}}(T)$ which is shown in Fig. SI: 5. In the range of integration time, $T \in [1, 2500]$, 64 input levels is adequate to accurately estimate $c_{\text{ideal}}(T)$, because increasing the number of input values to 128 produces no noticeable difference (less than 0.04 bits). We recommend always testing the estimate of $c_{\text{ideal}}(T)$ for multiple number of input levels $z$, to assure that $z$ is sufficiently large to avoid underestimation of $c_{\text{ideal}}(T)$.

### SI: 3. STOCHASTIC SIMULATIONS OF CENTRAL DOGMA MASTER EQUATIONS

#### A. Parameters for Fig. 2(a) and 2(b)

To produce the information gain curves in Fig.2(a) and 2(b), the following parameters were chosen to model the central dogma system, leakiness, $l = 0.01$, frequency parameter $\alpha = 1.0\,\text{min}^{-1}$, and input values $X \in [0, 1]$, which determines $k_{ON}$ and $k_{OFF}$ using Eq. (6). For the ideal information gain curves, $c_{\text{ideal}}(T)$, in Fig. 2(a), the transcript decay rate was $k_{d,m} = 0.5\,\text{min}^{-1}$, and the transcription rate constant $k_m = \{0.125, 0.5, 2.0, 8.0\}\text{min}^{-1}$ was determined using the transcription power $k_m/k_{d,m}$ values shown in Fig. 2(a). The set of integration times was $T = \{1, 2, 5, 10, 25, 50, 100, 250, 500, 1000, 2500\}$. The distribution of the ideal integration output was obtained using Eq. (21), which was then used to compute the ideal channel capacity, $c_{\text{ideal}}(T)$.

To produce the protein-level information gain curves in Fig. 2(b) the following central dogma rate constants were used: $k_m = k_{d,m} = 0.5\,\text{min}^{-1}$, the protein decay rate was $k_{d,g} = \{0.5, 0.2, 0.1, 0.05, 0.02, 0.01, 0.005, 0.002, 0.001, 0.0005, 0.0002\}\,\text{min}^{-1}$, and the translation rate constant was determined using the value of the translation power, $k_g/k_{d,g} =$

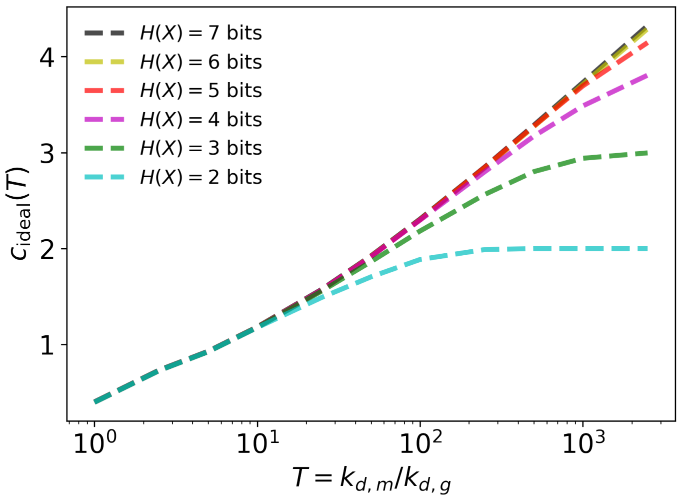


FIG. SI: 5. Ideal information gain $c_{\text{ideal}}(T)$ for increasing number of input values. Each ideal information gain curve was computed from $P(g_{\text{ideal}}^{(\text{analyt.})}|X)$ for $2^{H(X)}$ uniformly-spaced values of the input $X \in [0, 1]$.

$\{1, 10, 10^2, 10^3, 10^4\}$. For the translation powers $k_g/k_{d,g} \geq 10^2$ and integration time $T \geq 50$, we chose 6 bits sized input or 64 uniformly-spaced values of $X$ in $[0, 1]$. For all other translation powers and integration times, we chose 11 uniformly-spaced values of $X$ in $[0, 1]$. We performed Gillespie simulation of the central dogma master equations (7), (9), and (10) to obtain the protein expression distribution for each input value, $P(g|X)$, which was then used to compute the channel capacity, $c(X; g)$. Each stochastic simulation was run for a duration of $10^6$ min and $10^5$ samples of the protein expression level $g$ was obtained at a time interval of 10 min. The conditional distributions $P(g|X)$ were obtained from empirical distributions by binning the protein expression data. The number of bins, $n_b$, used were:

$$n_b = \begin{cases} 8 & \text{if} \quad T < 10 \\ 16 & \text{if} \quad 10 \leq T < 100 \\ 32 & \text{if} \quad 100 \leq T < 1000 \\ 64 & \text{if} \quad 1000 \leq T \end{cases} \tag{23}$$

#### B. Parameters for the four species, for Fig. 2(c) and 2(d)

To determine the ideal and the protein-level information gain for the four species we performed stochastic simulation of the central dogma reactions, equations (7), (9), and (10), using the central dogma rate constants in Table SI: 1.

| species | $k_m$ (min$^{-1}$) | $k_{d,m}$ (min$^{-1}$) | $k_g$ (min$^{-1}$) | $k_{d,g}$ (min$^{-1}$) |
|---|---|---|---|---|
| *E. coli* | 0.02 | 0.15 | $\{18, 9, 4.5, 1.8\}$ | $\{0.03, 0.015, 0.0075, 0.003\}$ |
| *S. cerevisiae* | 0.31 | 0.1 | $\{40, 20, 8, 4, 2, 0.8\}$ | $\{0.1, 0.05, 0.02, 0.01, 0.005, 0.002\}$ |
| *M. musculus* | 0.02 | 0.002 | $\{50, 20, 10, 5, 2\}$ | $\{0.002, 0.001, 0.0004, 0.0002, 0.0001\}$ |
| *H. sapiens* | 0.01 | 0.002 | $\{20, 10, 4, 2\}$ | $\{0.001, 0.0005, 0.0002, 0.0001\}$ |

TABLE SI: 1. Central dogma rate constants used to compute the typical protein-level information gain curves for the four species, solid lines with markers in Fig. 2(d). The translation loss region in Fig. 2(d) was computed from the ideal and the protein level information gain curves for integration time range $T \in [1, 1000]$.

The transcription and the translation rate constants were obtained from [9]. The transcript and the protein decay rate constants for each species were obtained from:

1. *E. coli* – Transcript decay rate constants from [10] and protein decay rates from [11].

2. *S. cerevisiae* – Transcript decay rate constants from [12] and protein decay rate constants from [13].

3. *M. musculus* – Both the transcript and the protein decay rate constants from [14].

4. *H. sapiens* – Transcript decay rate constants from [15] and protein decay rate constants from [16].

We determined the distribution of the dimensionless integration time, $T = k_{d,m}/k_{d,g}$, from the paired transcript and protein decay rate constants for each species (Fig. SI: 6). The distribution of integration times are shown as violin plots in Fig. 2(c). The 5% to 95% confidence interval was computed using numpy's percentile function in python, which ranks the samples and determines the percentile value using linear interpolation. The percentile values for the integration time, $T$, reported in the main text have been rounded off to the nearest integer. The violin plots in the main text were determined using matplotlib library's violinplot function with the 'scott' method for density estimate bandwidth.

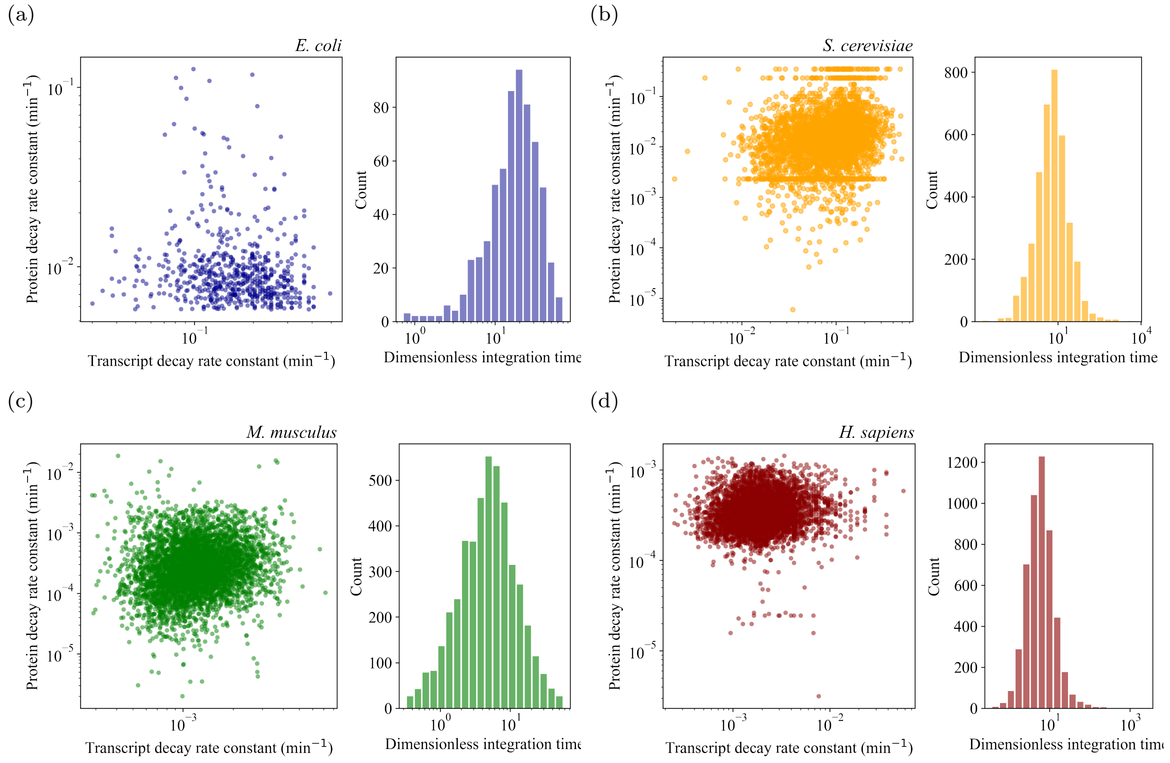


FIG. SI: 6. Paired transcript and protein decay rates and the distribution of dimensionless integration times for the four species. (a) Integration time distribution for *E. coli*. The effective protein decay rate was determined using a growth rate of $1/30\,\text{min}^{-1}$. (b) Integration time distribution for *S. cerevisiae*. (c) Integration time distribution for *M. musculus*. (d) Integration time distribution for *H. sapiens*.

The number of input levels, or values of $X$ in $[0, 1]$ for the Gillespie simulations, was chosen based on the integration time. Hence, for higher integration times a larger set of protein distributions, $P(g|X)$ was obtained from stochastic simulations to compute $c(X; g)$. The number of input values $X$ was selected by uniformly dividing the domain $[0, 1]$ into $2^{H(X)}$ intervals as,

For *E. coli* and *S. cerevisiae*:

$$H(X) = \begin{cases} 4 & \text{if} \quad T \leq 10 \\ 5 & \text{if} \quad 10 < T \leq 200 \\ 6 & \text{if} \quad 200 < T \leq 1000 \end{cases} \tag{24}$$

For *M. musculus* and *H. sapiens*:

$$H(X) = \begin{cases} 4 & \text{if} \quad T \le 10 \\ 5 & \text{if} \quad 10 < T \le 40 \\ 6 & \text{if} \quad 40 < T \le 200 \\ 7 & \text{if} \quad 200 < T \le 1000 \end{cases} \tag{25}$$

Gillespie simulations for each $X$ were mainly performed for $10^6$ min with the protein expression value sampled at an interval of 10 min to obtain $10^5$ samples of the protein expression level $g$. Except for *M. musculus* and *H. sapiens* when $50 < T \le 200$ the sampling interval was 100 min, and when $500 \le T$ the sampling interval was 200 min.

The protein expression distributions $P(g|X)$ were determined as the empirical distributions from the protein expression trajectory $g(t)$. The number of bins $n_b$ used to construct the empirical distributions were chosen as $n_b$ = nearest integer larger than $2^{c_{\text{ideal}}(T)+\eta}$, where $\eta > 0$, to use a larger number of bins than $2^{c_{\text{ideal}}(T)}$. For the three eukaryotic species:

$$\eta = \begin{cases} 2.0 & \text{if} \quad c_{\text{ideal}}(T) \le 1.0 \\ 1.5 & \text{if} \quad 1.0 < c_{\text{ideal}}(T) \le 2.0 \\ 1.0 & \text{if} \quad 2.0 < c_{\text{ideal}}(T) \end{cases} \tag{26}$$

and for *E. coli*:

$$\eta = \begin{cases} 2.5 & \text{if} \quad c_{\text{ideal}}(T) \le 0.1 \\ 3.0 & \text{if} \quad 0.1 < c_{\text{ideal}}(T) \le 1.0 \\ 2.0 & \text{if} \quad 1.0 < c_{\text{ideal}}(T) \le 1.5 \\ 1.5 & \text{if} \quad 1.5 < c_{\text{ideal}}(T) \end{cases} \tag{27}$$

For a more elaborate discussion on the selection of number of bins for computing channel capacity check [2–4].

#### C. Effect of fluctuation time period on information transfer

To determine how the time period of fluctuation of the input affects the information transfer from the input to the protein expression level, we determined the mutual information between the input and the ideal integration output $I(X; g_{\text{ideal}})$ under fluctuating protocols of the input, $X$. We chose the same $\alpha, l, k_m$, and $k_{d,m}$ used in the simulation study for Fig. SI: 4, and selected two values of the protein decay rate constant $k_{d,g} = k_{d,m}/10$ and $k_{d,m}/100$, which has integration times $T = 10$ and 100, respectively. For each of those two integration times we computed the ideal channel capacity $c_{\text{ideal}}(T)$ and the associated optimal input distribution, $P_{\text{opt}}(X)$ – the input distribution that achieves the channel capacity (Fig. SI: 8a). Then we considered a range of fluctuation time period for the input, $\tau_X = \frac{1}{k_{d,g}}\{0.1, 0.2, 0.5, 1, 2, 5, 10, 20, 50, 100, 200\}$. For each $\tau_X$ we performed a Gillespie simulation to capture the transcript trajectory $m(t)$ when the input $X$ fluctuates with time period $\tau_X$ assuming values according to the distribution $P_{\text{opt}}(X)$, the total duration of each simulation was $10^4\tau_X$. The stochastic transcript trajectory, $m(t)$, was then convoluted with the integration kernel $e^{-k_{d,g}t}$ with $t \in [0, 4/k_{d,g}]$, to obtain $g_{\text{ideal}}(t)$, which was subsequently used to compute the mutual information $I(X; g_{\text{ideal}})$. An example of the stochastic trajectories under a fluctuating protocol of the input, $X$, is shown in Fig. SI: 7.

For the two integration times $T = 10$ and 100, we obtained a set of ideal mutual information values, $I(X; g_{\text{ideal}})$, as a function of the fluctuation time period of the input, shown in Fig. SI: 8b. We notice in Fig. SI: 8b, when the time period of fluctuation $\tau_X$ is larger than translation response time $1/k_{d,g}$, almost by a factor of 10, then the ideal mutual information value $I(X; g_{\text{ideal}})$ approaches the ideal channel capacity for that integration time. When the fluctuation time period is smaller than $5/k_{d,g}$, then the ideal mutual information is less than half of the ideal channel capacity value. So, a relatively slow fluctuation in the environmental input is necessary to achieve the information gain possible due to integration of the transcript expression.

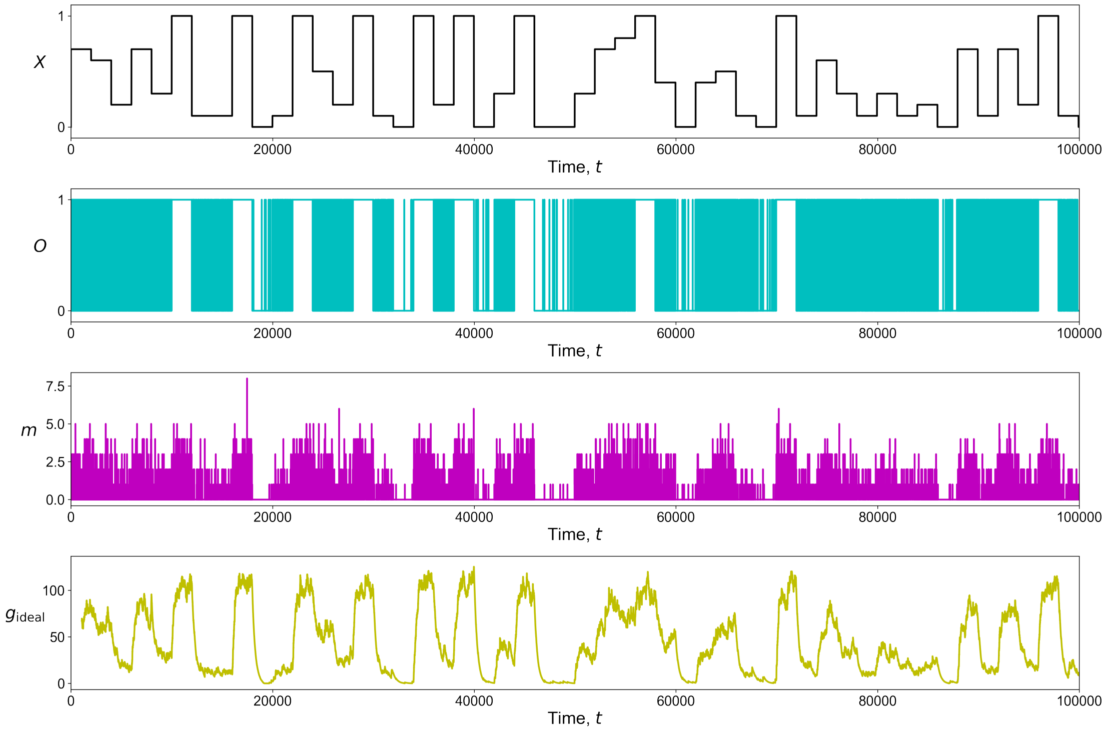


FIG. SI: 7. An example stochastic trajectory with the operator state $O$, transcript expression level $m$, and the ideal integration output $g_{\mathrm{ideal}}$, under a fluctuating protocol of the input, $X$. The input $X$ is changed stochastically after an interval of $\tau_X$.

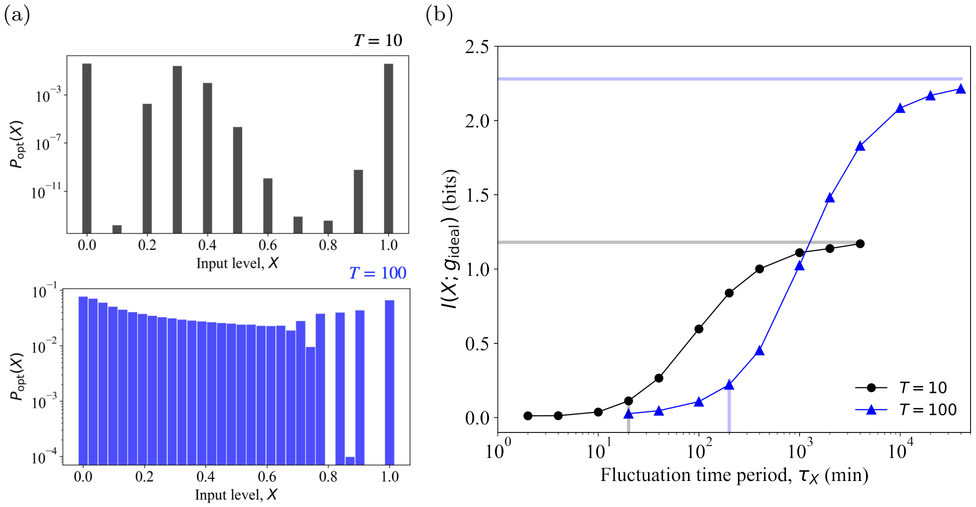


FIG. SI: 8. Ideal protein-level mutual information under a fluctuating protocol of the input, with fluctuation time period $\tau_X$. (a) Optimal input distribution, $P_{opt}(X)$, that achieves the ideal channel capacity, $c_{\mathrm{ideal}}(T)$, for integration times of 10 and 100. (b) Mutual information between the input $X$ and the ideal integration output $g_{\mathrm{ideal}}$ for fluctuating protocols with the same input distribution $P_{\mathrm{opt}}(X)$ but with different fluctuation time periods, $\tau_X$. The horizontal translucent lines represent the channel capacity $c_{\mathrm{ideal}}(T)$. $k_{d,m} = 0.5\,\mathrm{min}^{-1}$ for both the cases, so $k_{d,g} = 0.05\,\mathrm{min}^{-1}$ for $T = 10$ and $k_{d,g} = 0.005\,\mathrm{min}^{-1}$ for $T = 100$, respectively. The vertical translucent lines represent the translation response times $1/k_{d,g}$.

[1] R. Blahut, Computation of channel capacity and rate-distortion functions, IEEE transactions on Information Theory **18**, 460 (1972).
[2] R. Cheong, A. Rhee, C. J. Wang, I. Nemenman, and A. Levchenko, Information transduction capacity of noisy biochemical

signaling networks, science **334**, 354 (2011).

[3] S. Sarkar, D. Tack, and D. Ross, Sparse estimation of mutual information landscapes quantifies information transmission through cellular biochemical reaction networks, Communications Biology **3**, 1 (2020).

[4] R. Suderman, J. A. Bachman, A. Smith, P. K. Sorger, and E. J. Deeds, Fundamental trade-offs between information flow in single cells and cellular populations, Proceedings of the National Academy of Sciences **114**, 5755 (2017).

[5] J. Rammohan, Results of single-cell and single-transcript measurements collected for bias and resolvability attribution using split samples (brass) study, National Institute of Standards and Technology (2020), https://doi.org/10.18434/mds2-2300 (Accessed 2021-08-30).

[6] J. Rammohan, S. Lund, N. Alperovich, V. Paralanov, E. Strychalski, and D. Ross, Comparison of bias and resolvability in single-cell and single-transcript methods, Submitted to Communications Biology (2020).
